# Expression of a Barhl1a reporter in subsets of retinal ganglion cells and commissural neurons of the developing zebrafish brain

**DOI:** 10.1101/2020.02.20.957795

**Authors:** Shahad Albadri, Olivier Armant, Tairi Aljand-Geschwill, Filippo Del Bene, Matthias Carl, Uwe Strähle, Lucia Poggi

**Affiliations:** Centre for Organismal Studies, University of Heidelberg, Heidelberg, Germany; Institute of Biological and Chemical Systems, Biological Information Processing Karlsruhe Institute of Technology, Eggenstein-Leopoldshafen, Germany; Sorbonne Université, INSERM, CNRS, Institut de la Vision, Paris, France; Laboratory of Translational Neurogenetics, Department of Cellular, Computational and Integrative Biology - CIBIO, University of Trento, Trento, Italy; Laboratory of Molecular and Cellular Ophthalmology, Department of Cellular, Computational and Integrative Biology - CIBIO, University of Trento, Trento, Italy

**Keywords:** Barhl1a_1_, Barhl genes_2_, retinal ganglion cells_3_, retinal cell lineage_4_, neuronal subtypes identity_5_, Atoh7_6_, optic chiasm_7_, post-optic commissure_8_

## Abstract

Promoting the regeneration or survival of retinal ganglion cells (RGCs) is one focus of regenerative medicine. Homeobox Barhl transcription factors might be instrumental in these processes. In mammals, only *barhl2* is expressed in the retina and is required for both subtype identity acquisition of amacrine cells and for the survival of RGCs downstream of Atoh7, a transcription factor necessary for RGC genesis. The underlying mechanisms of this dual role of Barhl2 in mammals have remained elusive. Whole genome duplication in the teleost lineage generated the *barhl1a* and *barhl2* paralogues. In the Zebrafish retina, Barhl2 functions as determinant of subsets of amacrine cells lineally related to RGCs independently of Atoh7. In contrast, *barhl1a* expression depends on Atoh7 but its expression dynamics and function have not been studied. Here we describe for the first time a Barhl1a:GFP reporter line *in vivo* showing that Barhl1a turns on exclusively in subsets of RGCs and their post-mitotic precursors. We also show transient expression of Barhl1a:GFP in diencephalic neurons extending their axonal projections as part of the post-optic commissure, at the time of optic chiasm formation. This work sets the ground for future studies on RGC subtype identity, axonal projections and genetic specification of Barhl1a-positive RGCs and commissural neurons.

## Introduction

BarH-like homeodomain (BARHL) transcription factors play crucial roles in the control of neural cell fate specification, migration, subtype identity acquisition and survival during development of the retina and the brain ^1–7^. Studies have also implicated Barhl in neurodegenerative and neoplastic disorders ^8,9^. In *Xenopus* and mouse, *barhl2* appears to be the only family member expressed in the retina ^10,11^. In tetrapods, *barhl2* (previously named *MBH1* and *XBH1*) is expressed in both amacrine cells and RGCs of the developing and mature retina ^12,13^. Studies have also reported that Barhl2 is both sufficient and essential for determining the subtype specific identity of amacrine cells as well as to promote the maturation and survival of RGCs downstream of Atoh7 (also known as Ath5) – a bHLH transcription factor required for the specification of RGCs in vertebrates ^12–18^.

In Zebrafish, two *barhl* members, *barhl1a* (previously named *barhl1.2*) and *barhl2*, are expressed in the retina ^4,19^. We also have previously shown that, similarly to the mammalian counterpart, Barhl2 is an amacrine cell subtype identity-biasing factor downstream of the transcription factor Ptf1a ^20^. Furthermore, we took advantage of the zebrafish transgenesis combined with accessibility to 3D time-lapse imaging ^21–25^ to resolve the cell lineage origin of *barhl2*-expressing amacrine subtypes. Our study revealed that *barhl2*-expressing amacrine subtypes consistently arise within the lineage of Atoh7 upon reproducible asymmetrical divisions of RGC progenitors ^20^ (see also Fig. 8).

**Figure 1.**
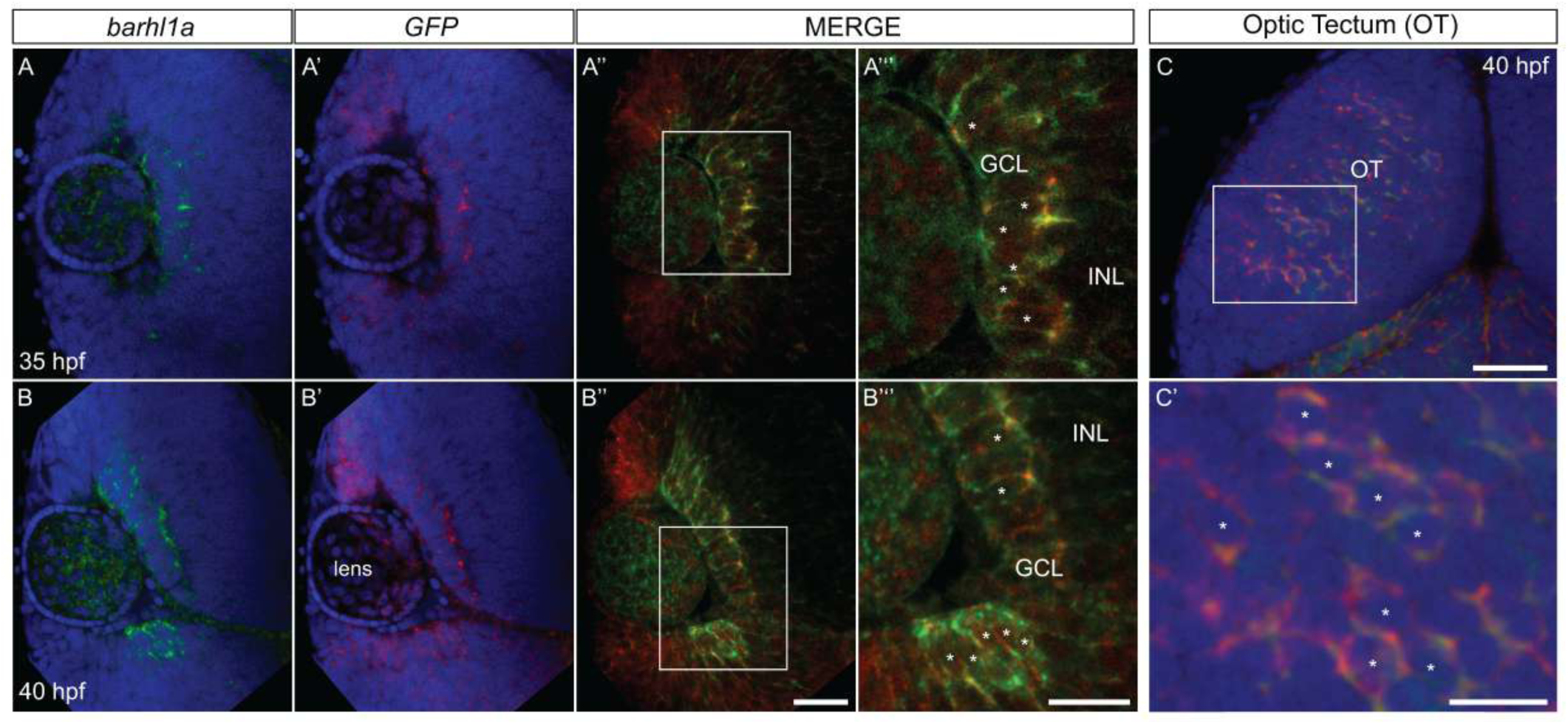
Endogenous *barhl1a* expression is faithfully reflected by the Tg(*barhl1a:GFP*) BAC-transgenic line in the retina and brain. Frontal optical sections of double fluorescent *in situ* hybridization (FISH) against endogenous *barhl1a* and *GFP* transgene in the retina and the optic tectum at 35 and 40 hours post-fertilization (hpf). (**A, B**) For both analyzed stages, *barhl1a* transcripts were found localized in the ganglion cell layer (GCL) as previously reported. (**A** – **B”’**) Faithfully reflecting *barhl1a* mRNA distribution, the *GFP* transcripts were also found restricted to the GCL layer co-localizing with *barhl1a* transcripts at both developmental stages. (**C** – **C’**) The same was observed in the tectal brain region where *barhl1a* expression was found overlapping with the one of the *GFP* transgene. GCL: ganglion cell layer, INL: inner nuclear layer, OT: optic tectum. Scale bars: (A – A”, B – B”, C) 50 *µ*m, (A”’, B”’, C’) 20 *µ*m.

**Figure 2.**
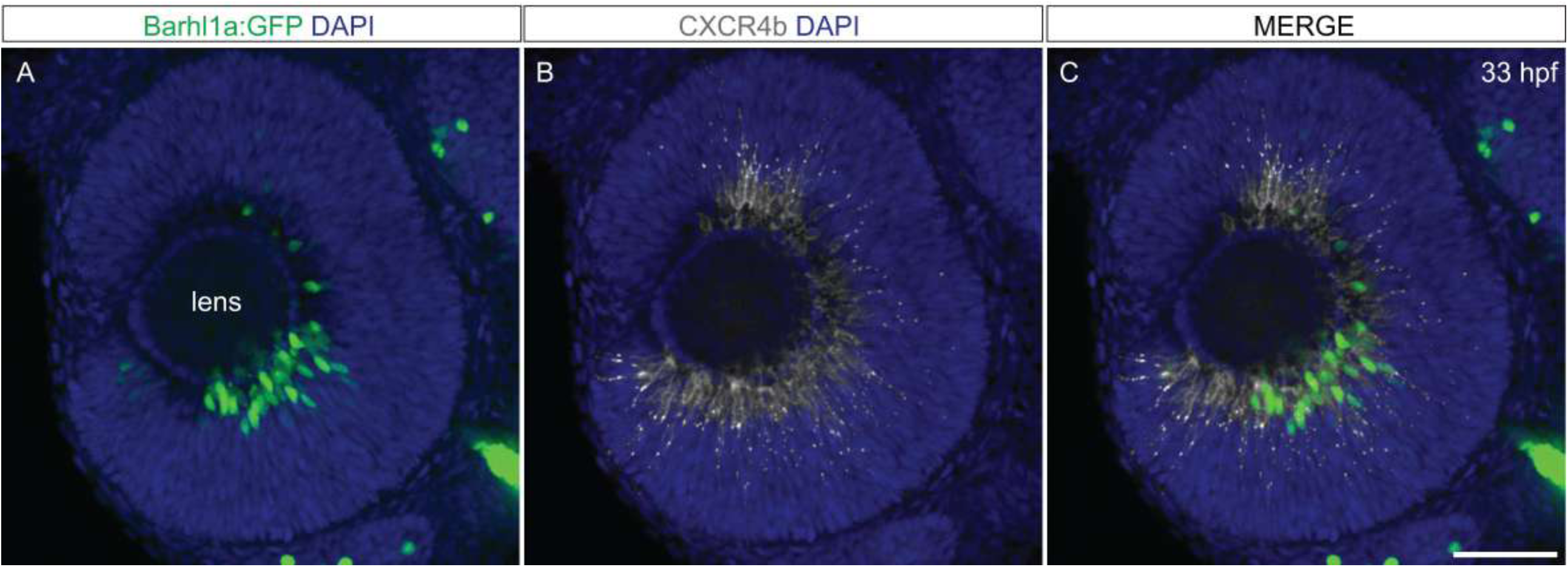
Tg(*barhl1a:GFP*) labels Cxcr4b post-mitotic retinal ganglion cells. Lateral optical sections of immunolabeled 33 hours post-fertilization (hpf) Tg(*barhl1a:GFP*) embryos with anti-Cxcr4b antibody. (**A**) GFP is detected in cells located basally around the lense at 33 hpf, colabeling with Cxcr4b (**B** and **C**), marking post-mitotic retinal ganglion cells and precursors. Scale bar: (C) 50 *µ*m.

**Figure 3.**
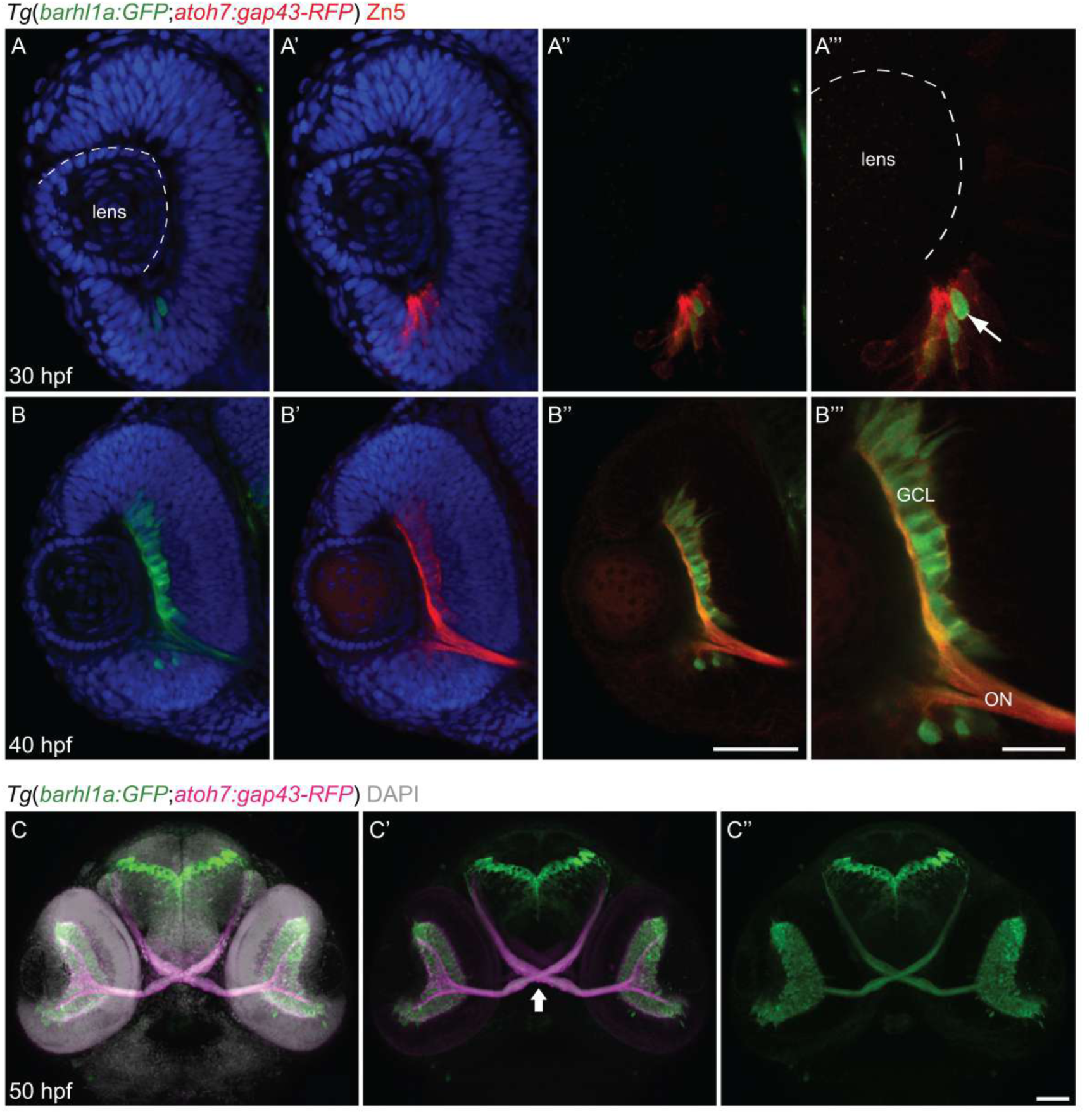
Barhl1a is expressed in a subset of *atoh7*-expressing retinal ganglion cells. Frontal optical sections of double transgenic Tg(*barhl1a:GFP; atoh7:gap43-RFP*) embryos at 30, 40 and 50 hours post-fertilization (hpf) counterstained with DAPI. (**A** - **A”’**) Barhl1a:GFP cells are detected at 30 hpf within atoh7:RFP-positive retinal ganglion cells at the ventro-nasal patch in the retina (while arrow, A”’). (**B** - **B”’**) At 40 hpf, GFP and RFP -positive cells in the ganglion cell layer (GCL) have extended their axons out of the retina to form the optic nerve (ON). (**C** - **C”**) By 50 hpf, Barhl1a-GFP cells have fully differentiated and extended their axons out of the retina forming the optic nerve. The optic nerves cross contra-laterally to form the optic chiasm (white arrowhead, **C’**) and reach their targets in the brain. At this developmental stage, no GFP could be detected from the pre-optic diencephalon. Scale bars: (A - A”, B - B”, C - C”) 50 *µ*m, (A”’, B”’) 20 *µ*m.

**Figure 4.**
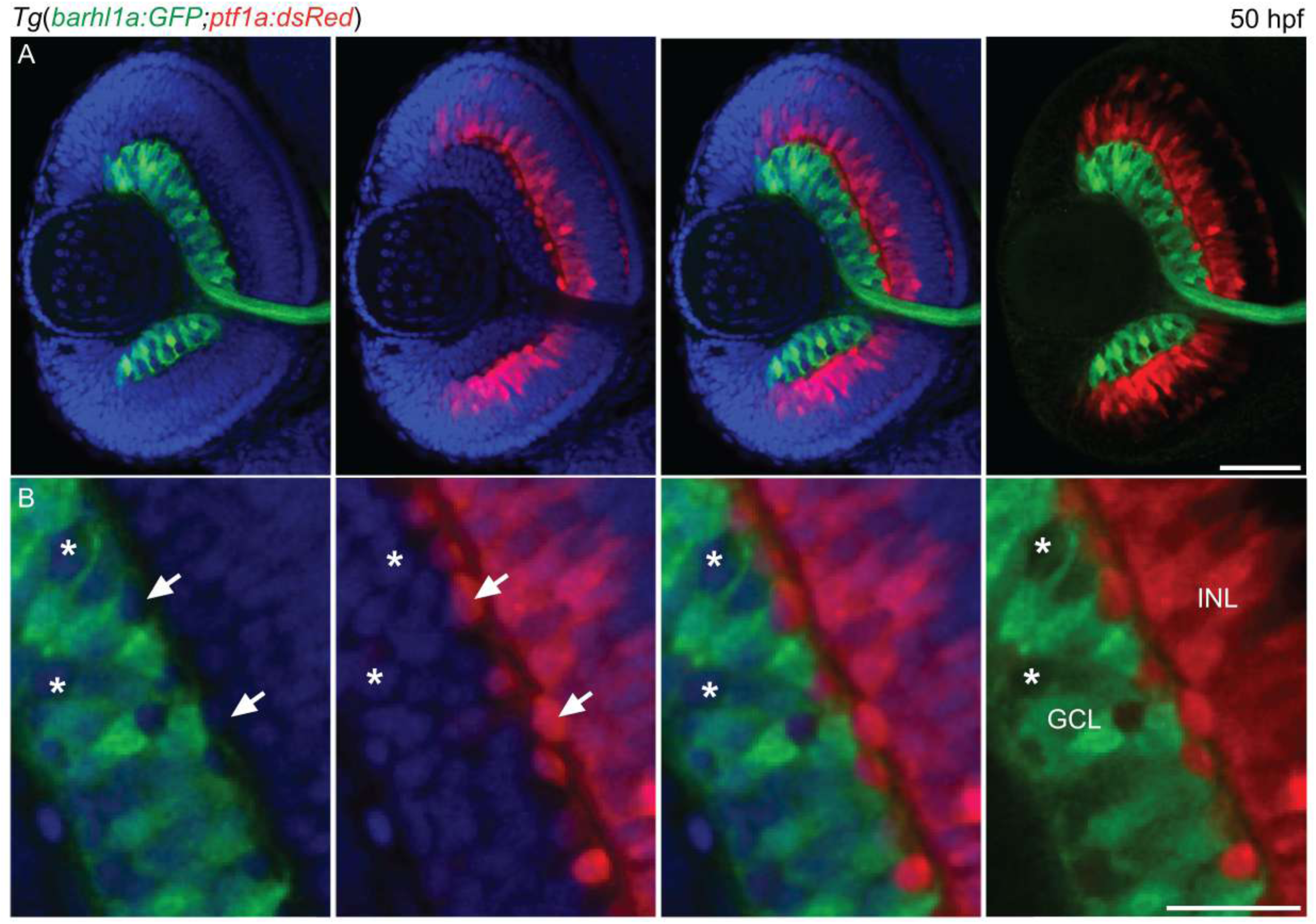
Barhl1a:GFP in the ganglion cell layer solely labels retinal ganglion cells. Frontal optical sections of double transgenic Tg(*barhl1a:GFP; ptf1a:dsRed*) embryos at 50 hours post-fertilization (hpf) counterstained with DAPI. (**A**) Barhl1a-GFP cells are detected in the ganglion cell layer (GCL) while DsRed-positive cells are detected in the GCL, in the inner nuclear layer and outer nuclear layer, marking displaced amacrine cells (white arrows, **B**), amacrine cells and horizontal cells. GFP-positive cells were never found co-labeling with DsRed, indicating that Barhl1a only marks retinal ganglion cells. Within the GCL, not all cells are Barhl1a:GFP-positive (white asterisks, **B**). Scale bars: (A) 50 *µ*m, (B) 20 *µ*m.

**Figure 5.**
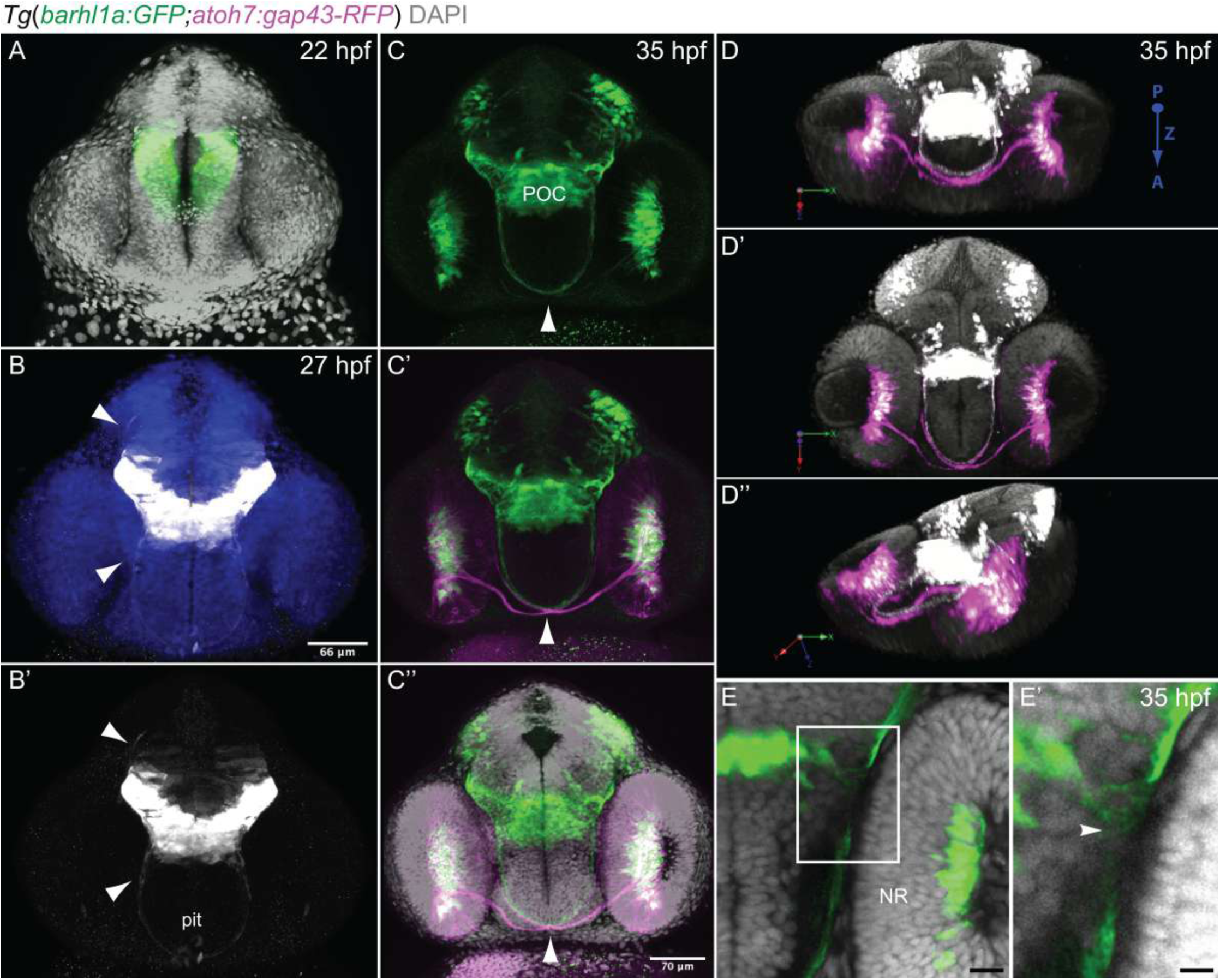
Developmental dynamic of Barhl1a-GFP cells in the diencephalon. Frontal optical sections and 3D reconstruction of transgenic Tg(*barhl1a:GFP;atoh7:gap43-RFP*) at 22, 27and 35 hours post-fertilization (hpf). (**A**) Barhl1a-GFP cells in the brain at 22 hpf are located in diencephalic domains around the midline and extend projections ventrally as observed at 27 hpf (white arrowheads, **B - B’**) to form a bundle of fibers along the optic tract. At this stage, few GFP-positive cells corresponding to pituitary cells could also be detected (pit, **B’**) By 35 hpf, retinal ganglion cells extended their axons out of the retina to form the optic nerve. The optic nerves extend from each eye, cross contra-laterally at the optic chiasm and extend along the optic tract in close proximity to the diencephalic GFP-positive fibers forming the post-optic commissure (POC) (white arrowhead, **C - C”**). (**D - D”**) 3D reconstruction of double transgenic Tg(*barhl1a:GFP;atoh7:gap43-RFP*) embryo at 35 hpf highlighting the growing optic nerves alongside GFP-positive fibers extending from the diencephalon (arrowheads, **E - E”**); in all three panels the view is from the posterior side of the stack/embryo; P, posterior; A, anterior; Z, z-axis. Scale bars: (A) 28 *µ*m, (B - B’) 66 *µ*m, (C - C”) 70 *µ*m, (E) 20 *µ*m, (E’) 50 *µ*m.

**Figure 6.**
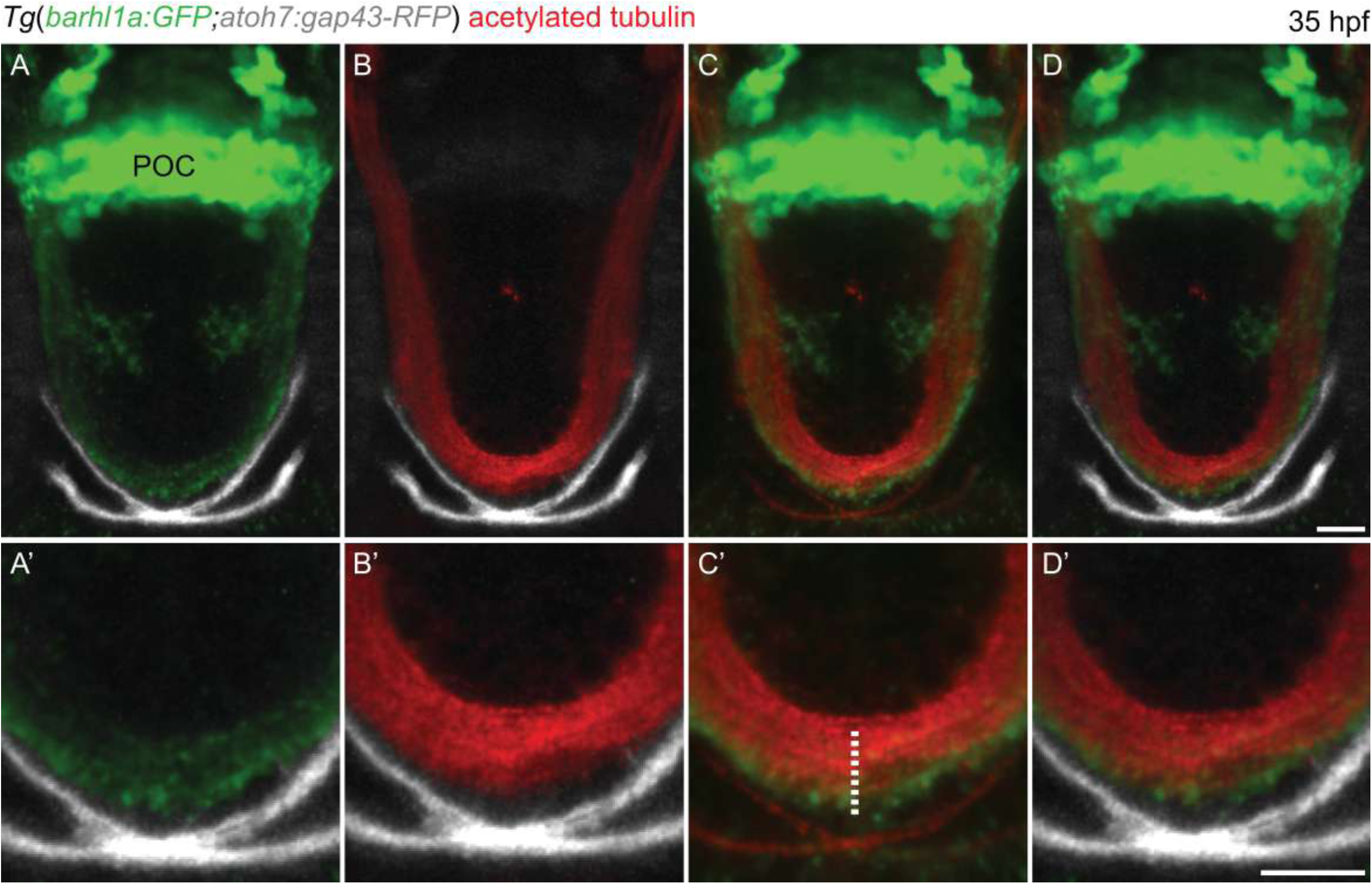
Barhl1a:GFP fibers partially co-label with the neuronal projection marker acetylated Tubulin. Acetylated-Tubulin immunostaining, labeling neuronal processes (**B - D’**), on Tg(*barhl1a:GFP;atoh7:gap43-RFP*) at 35 hours post-fertilization (hpf) reveals a partial overlap with the projections of Barhl1a:GFP cells marking the post-optic commissure (POC, **A**) in the diencephalon (overlap highlighted by the dash line in **C’**). Scale bars: (A - D) 50 *µ*m, (A’ - D’) 20 *µ*m.

**Figure 7.**
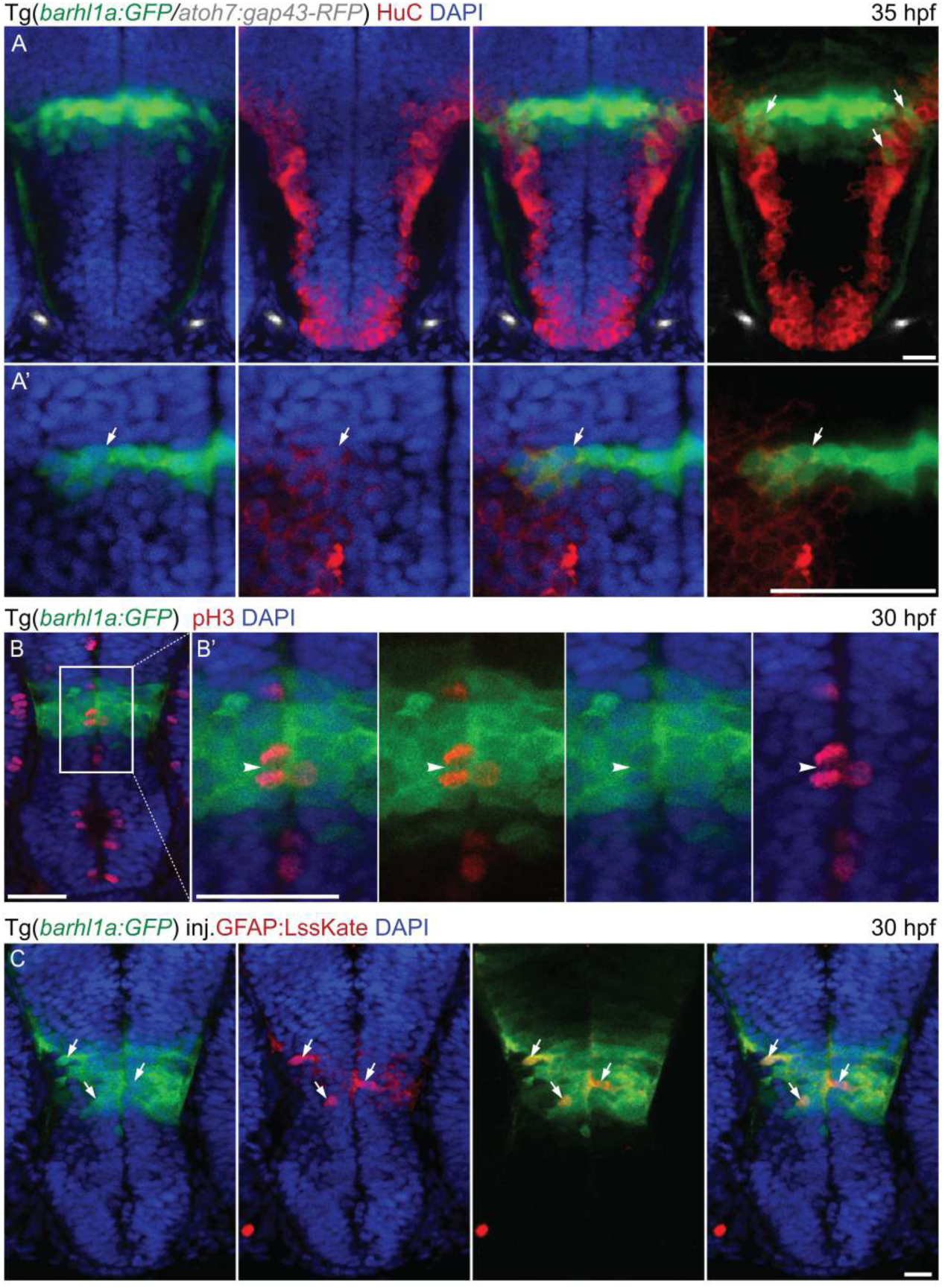
Barhl1a:GFP in the preoptic diencephalon marks populations of progenitor cells at the midline differentiating into neuron and glial cells. (**A - B**) Frontal micrographs of Tg(*barhl1a:GFP/atoh7:gap43-RFP*) embryos at 35 hours post-fertilization (hpf) immunohistochemically stained with anti-GFP antibody and anti-HUC (**A – A’**) and anti-GFP antibody and anti-pH3 (**B – B’**) antibodies. HuC marks the cell bodies of differentiated neurons and partially co-labels with Barhl1a:GFP cells at the margin of the GFP-positive preoptic diencephalon domain at 30 hpf (white arrows, **A - A’**). Close to the midline, pH3 labels cells dividing cells. PH3-positive cells could be detected within the domain of GFP-positive cells, (B and white arrowheads, **B’**). (**C**) Transiently injected Tg(*barhl1a:GFP*) embryos with the DNA construct *gfap:lssKate* labeling glial cells shows GFP and LssKate co-localization in the GFP-positive diencephalic domain at 30 hpf. Scale bars: (A, B, C) 50 *µ*m, (A’, B’) 20 *µ*m.

**Figure 8.**
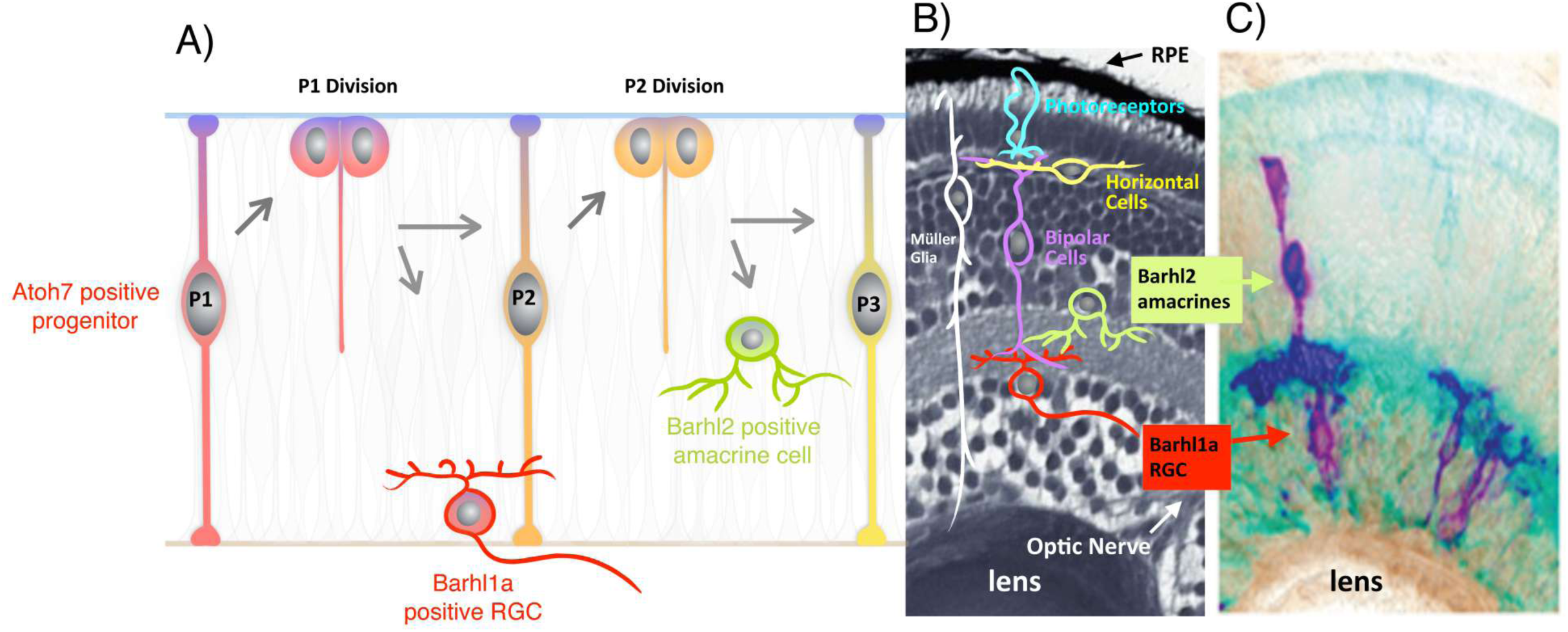
Asymmetric self-renewing division of Atoh7-expressing RGC progenitor giving rise to one Barhl1a-RGC and one Barhl2-amacrine cell. An hypothetical scenario is described, in which Barhl1a RGCs and Barhl2 amacrine cells tend to be clonally related and to become synaptic partners (see also ^20^).

Interestingly, while *barhl2* turns on exclusively in amacrine cells under the control of Ptf1a, the expression of *barhl1a* depends on Atoh7 and appears therefore to be restricted to the ganglion cell layer ^19^. The generation pattern of individual *barhl1a*-expressing cells within the Atoh7 cell lineage, the lineage relationship among Barhl1a and Barhl2 -expressing cells *in vivo* as well as the role played by Barhl1a in RGC genesis have so far remained unknown.

To address these questions, direct and dynamic visualization of *barhl1a*-expressing cells is needed. We used Tol2 transposase-mediated BAC transgenesis to generate a Barhl1a reporter line where GFP expression is driven by *barhl1a* regulatory regions. This reporter faithfully recapitulates *barhl1a* mRNA expression and helped us determine that Barhl1a turns on exclusively in a sub-population of RGCs after the cell cycle exit of *atoh7-*expressing progenitors. In addition, the reporter allows us to visualize axonal connectivity derived from *barhl1a-*expressing neurons. Furthermore, we provide the first description of the expression dynamic of Barhl1a in commissural neurons of the forebrain. This study thereby provides the foundation for further investigations of the role of Barhl1a transcription factors in nerve cell subtype identity acquisition and maintenance in this *in vivo* model system.

## Results

### Barhl1a in the retina is up-regulated in a subset of *atoh7*-expressing post-mitotic RGC precursors

In order to visualize *barhl1a*-expressing cells *in vivo* we generated a *Tg(barhl1a:GFP)* transgenic line expressing the reporter GFP under *barhl1a* regulatory genomic elements. To validate this transgenic line and to verify that the transcriptional activity of the *Tg(barhl1a:GFP)* transgene reflects the expression of the endogenous *barhl1a*, we first compared the spatial-temporal expression pattern of *barhl1a* mRNA and *GFP* mRNA in *Tg(barhl1a:GFP)* embryos by double fluorescent *in situ* hybridization (FISH). We found that expression of *GFP* in *Tg(barhl1a:GFP)* embryos reliably follows the spatial-temporal pattern of the endogenous *barhl1a* as previously described ^4,19,26^(**Fig. 1**). Transcripts of both genes can be consistently found as shown for example at 35 and 40 hours post fertilization (hpf), in the developing GCL as well as in the optic tectum (**Fig. 1**).

To directly investigate the dynamics of the appearance of individual Barhl1a:GFP-positive cells in the retina, we used 3D time-lapse imaging on *Tg(barhl1a:GFP)* embryos starting from 28 hpf. Imaging of the developing retina of *Tg(barhl1a:GFP)* embryos for at least 20 hours revealed that GFP positive cells become first visible in the ventral-nasal retina and spread subsequently to the whole retina, in a temporal and spatial fashion predictive of the wave of RGC differentiation (**Fig. 2 and Supplementary Movie 1**) ^27^. Individual cells turning on Barhl1a:GFP can be seen located either basally or in the apical half of the neuroepithelium to subsequently migrate basally until they reach the ganglion cell layer (**Fig. 2 and Supplementary Movie 1**). We never observed Barhl1a:GFP cells migrating to the apical surface to undergo division. This suggests that Barhl1a:GFP turns on in RGC precursors after the terminal division of *atoh7*-expressing neuroepithelial cells. Immuno-labeling of *Tg(barhl1a:GFP)* embryos with an antibody to Cxcr4b, a marker of post-mitotic RGCs ^28^ further supports our notion that Barhl1a:GFP is expressed in RGCs and their post-mitotic precursors (**Fig. 2**).

To investigate the dynamics of *Tg(barhl1a:GFP)* expression with respect to Atoh7 we generated a *Tg(barhl1a:GFP;atoh7:gap43-RFP)* double transgenic line in which the Atoh7:gap43-RFP labels all *atoh7*-expressing progenitors and their derived retinal cell types, including *barhl2-*expressing amacrine cells ^20,29^. Imaging the retina of this line shows that Atoh7:gap43-RFP becomes visible in the ventral-nasal retina already at 27-28 hpf, thereby highlighting the first-born RGCs ^27,30^. Two or three hours later, GFP expression from the Barhl1a reporter follows that of the Atoh7 reporter, being restricted to few Atoh7:gap43-RFP cells migrating to or residing within the GCL (**Fig. 3A-B”’**). At 50 hpf, expression of the Barhl1a reporter remains strong in the GCL and their axons have extended along the optic tract (**Fig. 3C-C”**). Of note, analysis of the 3D reconstruction of the optic chiasm and retinotectal projections of Barhl1a:GFP positive axons suggests that these projections reach specific tectal domains (**Supplementary Movie 2**).

Imaging in the retina of *Tg(barhl1a:GFP;atoh7:gap43-RFP)* embryos also revealed that not all Atoh7:gap43-RFP cells in the GCL appear as Barhl1a:GFP positive, as it would be expected if Barhl1a labels a subpopulation of RGCs (**Figs. 2, 4**). To assess this possibility, *Tg(barhl1a:GFP)* embryos were fixed and cryosectioned at 50 hpf. Sections where then immuno-stained against GFP and Zn5 (also known as Alcama/DM-GRASP/Neurolin/Zn8), a cell adhesion molecule that is transiently found on the entire surface of differentiated RGCs ^31^, and is therefore a suitable marker for labeling all RGCs. Barhl1a:GFP positive cell nuclei were counterstained with DAPI and imaging of the retinal sections was performed under a confocal microscope (see Materials). Quantification of the Barhl1a:GFP nuclei (1391 cells, n=6) in the central retina, where the retinal GCL is clearly distinguishable, shows that, on average, only about 54% of the entire RGC population is also Barhl1a:GFP positive.

Besides RGCs, the GCL also contains displaced amacrine cells. These displaced amacrine cells are however present in very low numbers, which would not largely affect the percentage of Barhl1a:GFP RGCs. We aimed nevertheless to rule out the possibility that some of the GFP-negative cells in the GCL may be displaced amacrine cells. To this end we generated a *Tg(barhl1a:GFP;ptf1a:RFP)* double transgenic line, in which the *ptf1a:RFP* transgene labels all amacrine and horizontal cells ^32–34^ (**Fig. 4**). Analysis of the retinae of *Tg(barhl1a:GFP;ptf1a:RFP)* embryos at 50 hpf revealed that expression of the two reporter genes are fully complementary, with all displaced amacrine cells expressing the Ptf1a reporter being Barhl1a:GFP-negative (**Fig. 4**). This further confirms that expression of *barhl1a* is restricted to a RGC population.

We conclude that Barhl1a turns on in about half the RGC precursors, which might mature at later stages during the wave of RGC genesis. The other half of the *atoh7*-expressing RGC precursors would then differentiate without the direct contribution of Barhl1a in the retina.

### Expression of the Barhl1a reporter in the brain labels diencephalic commissural neurons and their mitotic precursors

Prior to Barhl1a reporter activity in the retina, Barhl1a:GFP positive cells can be seen starting from 10-14 somites in the forebrain, in locations corresponding to the previously reported expression of *barhl1* mRNA in diencephalic domains of mammals, *Xenopus* and fish ^1,3,4,19,35^ (**Fig. 5A, B-B’ and Supplementary Movie 3**). These diencephalic domains have been shown to encompass the prospective pre-optic area, thalamus and posterior tuberculum/posterior hypothalamus ^1,3,4^. Subsequent GFP signals extend to more posterior domains, such as the ventral midbrain, pre-tectal/tectal area and the hindbrain ^3,4,36^ (**Fig. 5C-C” and Supplementary Movie 4)**. Finally, imaging of the forebrain of *Tg(barhl1a:GFP)* embryos additionally revealed previously not reported *barhl1a* expression in few cells of the pituitary (**Fig. 5B,B’ and Supplementary Movie S4**).

The Barhl1a reporter line offers the advantage to visualize not only the development of neuron cell bodies in the retina and brain but also of their axonal processes *in vivo* (**Fig. 5**). Imaging analysis of the brain of *Tg(barhl1a:GFP)* embryos indeed revealed, besides the optic chiasm, other two early commissural tracts deriving from *barhl1a* expressing cells. A first bundle of GFP positive fibers becomes visible at around 20-22 hpf, extending on both sides of the ventral-anterior diencephalon towards the midline, and forming a commissure around the presumptive optic tract (**Fig. 5B-B’ and Supplementary Movie 5**). The first RGCs become postmitotic in the retina only at around 27 hpf ^27^; we can therefore exclude that these Barhl1a:GFP-labelled axons crossing the ventral midline are RGC axons. Time-lapse imaging analysis using a lateral view further highlighted the temporal and spatial dynamics of the development of this ventral commissure in *Tg(barhl1a:GFP)* embryos, which appear before the optic chiasm and is located immediately posterior and dorsal to it (**Supplementary Movie 6 and see also Supplementary Movie 4**). Based on these data, we conclude that the Barhl1a:GFP positive ventral commissure is likely to contribute to the previously described post optic commissure (POC) ^37–40^. We then generated a BAC-based *barhl1a:gal4* (*barhl1.2:gal4-vp16*) DNA construct and injected it in one-cell stage *Tg*(*UAS:RFP;cry:GFP*) embryos, to obtain mosaic expression of RFP (**Supplementary Movie 7**). Single Barhl1a:gal4;UAS:RFP cells were seen extending their fibers along this POC; which are likely located in the caudal diencephalon/posterior tuberculum, where diencephalic commissural neurons have been previously described (**Supplementary Movie 7**) ^37–43^. We further found that Barhl1a:GFP overlaps with the expression of *lmx1b*, a gene expressed in posterior tubercular and hypothalamic neurons (**Supplementary Fig. S1**) ^44,45^. Remarkably, once the POC is fully formed and the optic nerves have reached their final destination, the Barhl1a:GFP commissural fibers are no longer visible (**Fig. 3C and Supplementary Movies 2, 6**). Therefore, the developmental spatial-temporal time of the appearance of the Barhl1a:GFP expression in the POC tightly correlates with the formation of the optic tract.

Another bundle of Barhl1a:GFP positive axons could be seen at around 27 hpf, which extend dorsally and cross the midline at the presumptive rostral border of the developing tectum (**Fig. 5B-C” and Supplementary Movie 6**). Based on their temporal appearance and anatomical position, these commissural fibers are likely to grow along the path pioneered by the posterior commissure (PC) and might be therefore residing in the ventral midbrain or pretectal nuclei (**Supplementary Movie 7**) ^37,39,43^.

To further investigate the location of the Barhl1a:GFP labeled diencephalic commissure with respect to the optic chiasm, we performed imaging in the brain of *Tg(barhl1a:GFP;atoh7:gap43-RFP)* embryos at 35 hpf (**Fig. 5 and Supplementary Movie S4**). At this developmental stage, Atoh7:gap43-RFP positive RGC axons can be clearly identified; which exited the retina and crossed the ventral midline to form the optic chiasm ^46^ (**Fig. 5C’-D”**). Conversely, only very few of these RFP -positive axons are also Barhl1a:GFP positive **(Fig. 5C - C”)**, further suggesting that the birth and/or maturation of Barhl1a:GFP RGCs is delayed with respect to the first wave of RGC genesis. Rotating the reconstructed 3D volume further highlighted the two contiguous but distinct commissures identified by the Atoh7 and Barhl1a reporter, respectively (**Fig. 5D-D”**). Lastly, *Tg(barhl1a:GFP;atoh7:gap43-RFP)* embryos were co-labeled immuno-histochemically with antibodies against acetylated Tubulin and GFP. The GFP signal overlaps with that of acetylated Tubulin in the POC fibers (**Fig. 6**), further indicating that the Barhl1a:GFP cell projections contribute to this diencephalic commissure.

In the pseudo-stratified neuroepithelium of the neural tube, post-mitotic neurons and glia are generated from progenitor cells that span the entire thickness of the neuroepithelium, from the basal to the apical surface. The nuclei of these progenitor cells undergo interkinetic nuclear migration, with mitotic nuclei being located at the apical surface whilst differentiating cells migrate to more basal locations ^47,48^. Since also Barhl1a:GFP positive cells span the entire thickness of the diencephalic neuroepithelium, we asked whether these cells would be found at different developmental phases of neurogenesis. *Tg(barhl1a:GFP)* neurons were firstly labeled immunohistochemically with a HuC/Elav antibody labeling the cell body of differentiated neurons ^49^. Overlap of HuC/Elav and GFP occurs in cells located further away from the apical (ventricular) surface, indicating that they are post-mitotic cells (**Fig. 7A**). Conversely, GFP positive cells immediately adjacent to the apical ventricular surface expressed the mitotic marker phospho-histone 3 (pH3) ^50^, indicating that they are mitotic precursors, as well as could be seen undergoing mitotic divisions (**Fig. 7B**). Neuronal progenitors in the zebrafish neuroepithelium have been reported to express markers of radial glia, such as the glial acidic fibrillary protein (GFAP) ^49,51^. Mosaic expression of an *gfap:lss-kate* construct, where the promoter of *gfap* has been cloned upstream of the *lss-mKate1* gene, highlighted GFAP/Barhl1a:GFP double positive cells at different apical-basal distances within the thickness of the neuroepithelium (**Fig. 7C**).

We conclude that, unlike in the neural retina, where *barhl1a* expression is turned on post-mitotically in RGC precursors, expression of the Barhl1a reporter in the diencephalon marks populations of progenitor/precursor cells undergoing cell division and neuronal and glial differentiation.

In summary, the observed spatial-temporal dynamics of Barhl1a:GFP are consistent with Barhl1a being a subtype identity factor for populations of commissural neurons in the brain as well as in the retina. The Barhl1a reporter indeed specifically labels particular neuronal subsets in the forebrain and their extending axons, which form commissures across brain hemispheres. These Barhl1a:GFP -expressing axons follow pre-existing embryonic commissures: the POC, the PC and the optic chiasm in a stereotypic pattern that is reminiscent of the previously described developmental sequence of the formation of archetypal tracts ^37,38,52,53^.

## Discussion

*Barhl* genes have a crucial role in the control of neuronal subtype acquisition and maintenance during development and in the adult ^1,2,5,6,8,14,54^. Loss of Barhl2 protein in the postnatal mammalian retina causes programmed cell death of more than 50% of RGCs ^13^. Therefore, a better understanding of Barhl function is essential to prevent RGC death and/or enable RGCs and axon regeneration. However, the mammalian Barhl2 is concomitantly expressed in amacrine cells and it is required for their subtype identity acquisition ^13,14^. Untangling these two concurrent functions of Barhl2 might therefore be challenging in mammals. Barhl proteins play evolutionarily conserved roles in retinal cell type maturation ^12,20^. Furthermore, the two zebrafish *barhl* paralogues genes, *barhl1a* and *barhl2*, exhibit complementary expression in the retina ^19^, which together resemble mammalian Barhl2 expression.

We here started to explore whether it could be possible to disentangle the dual function of the single mammalian Barhl2 gene in the retina, by analyzing its zebrafish orthologous genes. With the generation of the *Tg(barhl1a:GFP)* transgenic line we established an essential resource. This line allowed us to provide for the first time a comprehensive description of the dynamic *barhl1a* expression *in vivo*. Our study confirms and extends previous findings by showing that Barhl1a:GFP turns on exclusively in RGCs (most likely downstream of Atoh7; ^19^). Time-lapse imaging of individual Barhl1a:GFP -positive RGCs further enabled us to show that these cells are post-mitotic neuro-epithelial cells, with their nuclei migrating from the apical half of the neuroepithelium towards the GCL.

Interestingly, these Barhl1a:GFP RGCs represent only about 54% of the RGCs population, which appear to display distinct retinotectal projections. There is a delay in the appearance of Barhl1a:GFP positive axons compared to the appearance of Atoh7:gap43-RFP axons. One explanation for this delay might be that Barhl1a:GFP cells comprise a population of later-born RGCs, which grow their axons along the optic tract pioneered by the early-born Atoh7-positive/Barhl2-negative RGCs ^53^. Alternatively, the delayed onset of GFP could also reflect the requirement of Barhl1a expression for the late maturation and/or maintenance of RGCs. Conclusions cannot be drawn here and future studies using the *Tg(barhl1a:GFP)* line and *barhl1a*-inducible Gal4 constructs may be used to investigate the distinct molecular signature, morphology, dendritic patterns and retinotopic targets of Barhl1a RGCs.

We have previously shown that about 58% of amacrine cells express Barhl2 in the Zebrafish retina, all of which seem to arise from the asymmetric divisions of *atoh7*-expressing progenitors. Moreover, Zebrafish Barhl2 is both necessary and sufficient for subtype identity acquisition of these amacrine cells downstream of Ptf1a ^20^. Given the remarkably similar temporal and subtype-restricted expression pattern of Barhl1a, it will be interesting in future studies to examine if and how Barhl1a determines particular RGC subtype identities and retinotopic projections, and whether these subtypes tend to have recurrent clonal relationships and connectivity patterns with their lineally-related Barhl2 amacrine subtypes (**Fig. 8**). Lastly, the fact that Barhl1a RGCs and Barhl2 amacrine cells are lineally related (that is: they arise from the same type of progenitor) suggests that these retinal cells share gene expression profiles and/or epigenetic ancestry. It will be exciting to explore whether such insights gained from a genome duplication in zebrafish will allow to narrow down the shared and distinct gene networks that are relevant for the role of Barhl in the maturation of RGCs and amacrines as well as in the survival of RGCs in the mammalian retina. Understanding this could be instrumental, for instance, for the identification of key factors whereby reprogramming of lineally-related retinal cell types, such as Barhl-dependent amacrines and RGCs, could be achieved to prevent RGC death or encourage their regeneration from amacrine cells ^55^.

Unlike in the retina, where Barhl1a:GFP is restricted to post-mitotic neurons, Barhl1a:GFP in the brain is turned on in mitotic progenitors and retained in their post-mitotic daughters. Using time-lapse analysis in the brain of *Tg(barhl1a:GFP)* embryos we discovered for the first time that Barhl1a:GFP in the diencephalon labels small populations of cells contributing to the POC and the PC tracts. Intriguingly, we find partial overlapping expressions of Barhl1a:GFP and *lmx1b*, a regulator of dopaminergic neurons ^45^. Based on the mosaic expression of Barhl1a:GFP, we speculate that these POC neurons might be located in the basal diencephalon/posterior tuberculum, where they co-localize with *lmx1b*. Interestingly, the onset of Barhl1a:GFP labeled POC fibers coincides with the onset of RGC axon maturation and is transient, *i.e.* it disappears when the optic tract is fully differentiated. Future studies will assess the precise identity of Barhl1a:GFP commissural neurons; they will also assess whether the remarkably harmonized developmental pattern between the Barhl1a:GFP labeled POC and optic chiasm may underlie a molecular cross-talk required for the proper formation of the optic tract.

In conclusion, our novel experimental tools and insights into the spatial-temporal dynamics of Barhl1a:GFP in the retina and brain provide a fundamental framework for further investigation of Barhl1a-specific RGCs and their retinotectal RGC projections as well as of Barhl1a-commissural projection neurons *in vivo*. Such investigations are likely to be relevant to dissect BarHl functions in mammals and to understand and ameliorate the pathophysiology of BarHl-linked diseases.

## Methods

### Animals and ethic statement

Fish were maintained at 26-28°C and embryos raised at 28.5°C or 32°C and staged as described previously ^56,57^. Fish were housed in two facilities: Fish facility at COS, University of Heidelberg, Germany; fish facility at CIBIO, University of Trento, Italy. Each facility is under supervision of and in accordance with local animal welfare agencies and European Union animal welfare guidelines (Tierschutzgesetz 111, Abs. 1, Nr. 1; Regierungspräsidium Karlsruhe and the Italian Ministry of Health - permit no.: 151/2019-PR). Zebrafish (*Danio rerio*) embryos of either sex were used exclusively before free-feeding stages. Embryos used for whole-mount imaging were treated with 0.0045% 1-phenyl-2-thiourea (Sigma) to delay pigment formation. Lines used in this study were generated in the zebrafish wild type background (AB/AB or AB/WIK). The fish facility is under the supervision of the local representative of the Animal Welfare Agency.

### Fish lines

For the generation of the *Tg(barhl1a:GFP)* line, the BAC clone CH120-215H7 (CHORI, BACPAC, Oakland, CA, USA) was modified using the *flpe recombinase* system in EL250 cells ^58^. The coding sequence of eGFP with a SV40 polyA site was inserted within the first exon of the *barhl1.2* genomic sequence and the tol2-AmpR cassette was inserted in the TARBAC2.1 backbone by PCR. The modified BAC was injected together with Tol2 mRNA into one cell-stage zebrafish embryos to generate a stable transgenic line.

Along with the newly generated and validated Tg(*barhl1a:GFP*) line, five other transgenic lines expressing *GFP, dsRed* and *gap43-RFP* under the control of different promoters were used in this study: the previously published Tg(*atoh7:gap43-RFP*) ^29^; the derived outcrossed line Tg(*barhl1a*:*GFP;atoh7:gap43-RFP*); Tg(*barhl1a:GFP;ptf1a:dsRed*) generated by outcrossing Tg(*barhl1a:GFP*) and Tg(*ptf1a:dsRed*) line *Tg((−5.5ptf1a:DsRed)ia6)* previously published ^20^ and the previously published Tg(*UAS:RFP;cry:GFP*) transgenic line ^59^. Embryos carrying both transgenes were screened for expression of the red and GFP reporters using an Olympus MVX10 macrofluorescence binocular.

### Immunohistochemistry

The primary antibodies used in this study and their dilutions were the following: chicken anti-GFP antibody (Life Technologies, A10262, 1:500), rabbit anti-DsRed (Clontech, 632496, 1:200), mouse anti-Tubulin (Sigma T5168, 1:100), mouse anti-HUC (Molecular probes, A-21271, 1:100) rabbit anti-pH3 (Millipore, 06-570; 1:500), mouse anti-Zn5 (ZIRC Zn-5; 1:200), monoclonal zebrafish anti-Cxcr4b (1:100, ^60^. Secondary antibodies were goat or donkey anti-mouse, anti-rabbit or anti-goat IgG conjugated to Alexa Fluor 488, 546, 594 or 647 fluorophores (1:500 for whole-mount to 1:2000 dilutions for cryosections; Invitrogen).

Whole-mount immunohistochemistry experiments were carried out as follows: embryos were fixed 1 to 2 hours in 4% paraformaldehyde (4% PFA). The fixative was then washed out using 1X PBS-Tween solution. Permeabilization of the embryos was done on ice using 0.25% Trypsin-EDTA (with phenol red; Gibco, Life Technologies) solution. The duration of trypsin treatment was dependent on the stage of the embryos: 28–30 hpf:3 min; 31–33hpf:4min; 34–35hpf: 6min; 40hpf:7–8min; 45hpf: 9–10min; 48– 50hpf: 11–12 min; 60 hpf: 20 min. Embryos were then washed several times using 0.1 M PBS-Tween solution (1X PBS-Tween solution), blocked using 10% blocking solution (10% heat-inactivated goat serum, 1% bovine serum albumin, 0.2% Triton X-100 in PBS) and incubated overnight at 4 degrees in 1% blocking solution (10% blocking solution in 1X PBS-Tween) in which primary antibodies were diluted. Primary antibodies were then washed out using 1X PBS-Tween and embryos were then incubated overnight at 4 degrees in 1% blocking solution (1% heat-inactivated goat serum, 1% bovine serum albumin, 0.2% Triton X-100 in PBS) in which secondary antibodies and 4’,6-diamidino-2-phenylindole (DAPI) were diluted. Prior to imaging, embryos were washed in 1X PBS-Tween and stored at 4 degrees.

For the immunohistochemistry on sections, embryos were fixed in 4% paraformaldehyde (PFA) in 0.1 M phosphate saline buffer (PBS) overnight at 4°C, rinsed and cryoprotected in 30 % sucrose (w/v) overnight at 4°C. Embryos were then mounted vertically (head down) in freezing medium (Jung Tissue Freezing Medium, Leica Microsystems), frozen in liquid nitrogen and cryosectioned immediately with Leica CM3050 S cryostat. The thickness of sections was 14 µm. Sections were collected on adhesion microscope slides (SuperFrost Plus, Menzel-Gläser) and left to dry overnight at 4 °C. For immunohistochemistry, microscope slides with cryosections were washed 3 times (each wash 15 min) in PTW, then covered with 10 % goat blocking medium and incubated at room temperature for 1 h. The cryosections were then incubated overnight in primary antibodies: anti-Chicken GFP (Life Technologies, A10262) and anti-Mouse Zn5 (Zebrafish International Resource Center (ZIRC)) both diluted 1:500 in 1 % goat blocking medium. Microscope slides were then washed again 3 times (each wash 15 min) in PTW and secondary antibodies were added: anti-chicken conjugated to Alexa Fluor 488 (from donkey; Jackson ImmunoResearch Laboratories, Inc., 703-545-155) and anti-mouse conjugated to Alexa Fluor 546 (from goat; Invitrogen, A-11030) both diluted 1:500 in 1 % goat blocking medium. Cryosections were incubated with the secondary antibody mix for 1.5–2 h at 37 °C in the dark and then washed again in PTW (3 times 15 min) at room temperature. They were then incubated in DAPI (10 µg/ml in PTW) for 10–15 min and washed in PTW (3 times 15 min). Microscope slides were then dried from the back and edges with a paper tissue, 120 µl 60 % glycerol was added and cryosections were covered with coverslips (24×60 mm, Carl Roth). Coverslips were sealed with nail polish and microscope slides were stored at 4 °C in the dark until imaging.

### Double fluorescent *in situ* hybridization

For *barhl1a/GFP* double fluorescent whole mount *in situ* hybridization (FISH), standard digoxigenin- and fluorescein-labeled riboprobes were combined with Tyramide Signal Amplification, as described by ^19^. *Barhl1a* riboprobe was synthesized as previously reporter ^19^ and *gfp* riboprobe was synthesized from a linearized *pCS2:GFP* plasmid with NotI (Fermentas or New England Biolabs) and transcribed with Sp6 (mMessage mMachine Sp6, Ambion). Incubation with the *barhl1a* probe was for 40 minutes, incubation with the *gfp* probe was for 30 minutes. Embryos were then kept in the dark for all following steps. For detection and staining of the antisense probes, embryos were washed 5 times 10 minutes with TNT (0.1M Tris pH7.5, 0.15M NaCl, 0.1% Tween-20), incubated with 1% H2O2 in TNT for 20 min and washed again 5 times 10 minutes. Embryos were blocked in TNB (2% DIG Block in TNT) for 1 hour at room temperature and afterwards incubated with Anti-Digoxigenin-POD Fab fragments diluted 1:100 in TNB. For signal detection, Fluorescein (FITC), Cyanine 3 (Cy3) or Cyanine 5 (Cy5) Fluorophore Tyramide by PerkinElmer was used. Embryos were then incubated in 1X 4’,6-Diamidin-2-phenylindol (DAPI) in TNT overnight at 4°C and washed several times in TNT the next day. The stained embryos were then used for imaging or kept in the dark at 4°C until imaging.

For *barhl1a/lmx1b* double fluorescent whole mount FISH, antisense RNA probes for *barhl1a* and *lmx1b1* were produced from full length cDNA clones generated by ^61^. Antisense probes were generated as described in, ^61^ except that DNP-11-UTP (Perkin Elmer) was used for the production of *barhl1a* RNA probe. Fluorescent double whole mount FISH, RNA hybridization were performed on dechorionated 20 hpf embryos with the tyramide amplification kit (TSA Plus Cyanine 3 System, Perkin Elmer, Boston, MA). Briefly, manually dechorionated 20 hpf embryos were fixed in 4% (w/v) PAF for 3 hours at 4°C and conserved until use in 100% (v/v) Methanol at -20°C. Embryos were rehydrated through MetOH/PBS gradient series and washed 3 times in 0.1% (v/v) Tween 20 in PBS buffer (PTW). Endogenous peroxidase were inactivated in 2% (v/v) H_2_O_2_, the embryos incubated 2 minutes with proteinase K (10 µg/ml at room temperature (20°C), then postfixed 15 min in 4% PFA for 30 minutes at room temperature and washed 5 times 5 minutes in PBS-Tween. Embryos were then pre-hybridized 3 hours before overnight incubation at 65°C in hybridization buffer (pH 6) containing the antisense-labeled probes. On the second day, embryos were washed in post-hybridization buffers, incubated 1h in blocking buffer and then incubated overnight with an anti-DNP-POD antibody (1:1000, Perkin Elmer). The next day embryos were washed in PBS-Tween, and stained with tyramide Cy3 solution (1:100) in 0.002% (v/v) H_2_O_2_ in PBS-Tween, peroxidase were inactivated in 2% (v/v) H_2_O_2_ for 1 hour and washed in PBS-Tween. The same day, embryos were incubated overnight with an anti-digoxigenin-poly-POD antibody (1:1000, Roche), then washed in PTW and stained with tyramide Cy5 solution (1:100) in 0.002% (v/v) H_2_O_2_ in PBS-Tween.

### Constructs

The *barhl1a:gal4* bacterial artificial chromosome (BAC) construct was generated as follows. For the BAC manipulation, bacteria containing a BAC clone spanning the *barhl1a* genomic locus (CH211-215C18, BACPAC Resources Center) were used. Transformation through electroporation of the pRedET plasmid was performed as described (Bussmann and Schulte-Merker, 2011). For Tol2 transposon-mediated BAC transgenesis, the iTol2-amp cassette ^62^ was amplified by PCR with the primer pair pTarBAC_HA1_iTol2_fw (5’-gcgtaagcggggcacatttcattacctctttctccgcacccgacatagatCCCTGCTCGAGCCGGGCCCAAGTG-3’) and pTarBAC_HA1_iTol2_rev (5’-gcggggcatgactattggcgcgccggatcgatccttaattaagtctactaATTATGATCCTCTAGATCAG CTC-3’), where the lower and upper cases indicate the pTarBAC2.1 sequences for homologous recombination and the iTol2-amp annealing sequences, respectively. Subsequently, the amplified iTol2-amp cassette was introduced into the backbone (pTarBAC2.1) of the Barhl1a-BAC. 500 ng of the PCR product (1 mL) were used for electroporation. 5’-caaaaccagtgtcataaaggacaaatgcacatttgatattgatttgactcGCCACCATGAAGCTACTGTCTTCTATCGAAC-3’ and 5’-ctgtgagaaagtatagactcgatcccaaagctcgagccgtttgatacctcCCGCGTGTAGGCTGGAGCT GCTTC-3’ primers were used to amplify and insert the gal4FF cassette into the BAC ^63^. The lower and upper cases indicate the CH211-215C18 sequences for homologous recombination and the pGal4FF-FRT-Kan-FRT annealing sequences, respectively. 500 ng of the PCR product (1 mL) were used for electroporation in Barhl1a-iTol2-amp-BAC -containing cells. The BAC DNA preparation was performed using the HiPure Midiprep kit (Invitrogen), with modifications for BAC DNA isolation as described by the manufacturer. Tol2 transposase mRNA was prepared by in vitro transcription from XbaI-linearized pDB600 ^64^ using the T3 mMessage mMachine kit (Ambion). RNA was purified using the RNeasy purification kit (Qiagen), diluted to a final concentration of 100 ng/µl for injection.

The zebrafish *gfap* promoter ^51^ was derived from the Addgene plasmid: *pEGFP-gfap*(Intron1/5’/Exon1-zebrafish) (Addgene, #39761) and *lssmKate2* ^65^ was derived from the Addgene plasmid: pLSSmKate2-N1 (Addgene, #31867). In a first step the *gfap* promoter driving GFP was sub-cloned into a vector containing IsceI sites using NotI and XhoI restriction enzymes (NEB #R3189 and #R0146). To insert *lssmKate2* SalI and NotI restriction enzymes (NEB #R3138S and #R3189) were used, resulting in the following plasmid: *IsceI:: gfap(Intron1/5’/Econ1-zebrafish)::LssmKate2 polyA*.

### Imaging and statistical analysis

For either imaging of fixed embryos or live imaging, embryos were embedded in 1 % low-melting agarose (w/v, diluted in H2O; Biozym) in 35 mm Glass-bottom Microwell dishes (P35G-1.5-10-C, MatTek). They were oriented with truncated Microloader tips (Eppendorf) frontally (head touching the dish and body axis tilted about 45°) or laterally. After solidification, agarose was covered with a drop of water to avoid dehydration during imaging. Confocal microscopy was carried out with Leica SpE or Leica Sp5 confocal laser scanning microscope using a Leica 20x or 60x 1.2 NA water-immersion objective and Leica Application Suite (LAS) software. In most cases, confocal images were taken sequentially from the whole head/retina, with a distance of 2 to 5 µm in z-direction. Optical stack of 40–60 µm for time-lapse and a maximum of 100 µm for fixed embryos were imaged. Live imaging was performed as previously described ^21–23^. Images were taken every 5 or 10 minutes for 24 – 42 hours and the motorized XY stage was used to image multiple embryos. For all imaging experiments, laser power was minimized to avoid bleaching and photo-toxicity. Sequential image acquisition was also performed using individual descanned Leika PMT detectors. All image processing and 3D reconstructions were done using either Volocity 6.0.1 (PerkinElmer) or Fiji (https://imagej.net/ImageJ).

In order to quantify the number of Barhl1a:GFP cells in the GCL, cryosections immunostained with Zn5 were imaged with a Leica DM5000B compound microscope (20x objective). Three most central sections of each retina (n = 6 embryos) were imaged. Both GFP positive and GFP negative nuclei were counted manually in Volocity Analysis version 5.3 (Improvision). To exclude cells in the periphery, where the CMZ continues to give rise to all retinal cell classes, only cells within the central quadrant of the retina were counted as previously described ^20^. For every retina, the percentage of GFP-positive cells was calculated as a mean of the proportions observed in each individual section.

## Acknowledgments

We thank J. Wittbrodt and K. Lust for kindly providing the *gfap:lss-kate* DNA construct made by K. Lust in the laboratory of J.Wittbordt. We thank B. Wittbrodt, E. Leist, A. Saraceno, M. Majewski, B. Seiferling, T. Kellner at COS, University of Heidelberg, and I. Mazzeo and S. Robbiati at CIBIO, University of Trento, for fish maintenance and technical assistance. We are also thankful for the support of the core imaging facility at CIBIO, University of Trento. S.A. was supported by the Landesgraduiertenförderung (Funding program of the State of Baden Württemberg, Germany). This work was supported by Deutsche Forschungsgemeinschaft Research Grant PO 1440/1-1 to L.P and by University of Trento.

## Authors Contributions

L.P. conceptualized and supervised the project. S.A. performed the research and obtained and prepared figures. L.P. and S.A. wrote the manuscript together with M.C. F.D.B. U.S. commented and proofed the manuscript. M.C. contributed with the conceptualization and correction of the manuscript. S.A. generated the *Barhl1.2:gal4-vp16* BAC construct in the laboratory of F.D.B and performed mosaic expression experiments. O.A. and U.S. made the Tg(*barhl1a:GFP*) transgenic line and *in situ* analysis of *barhl1a* and *lmx1b*. T.A.G. performed Cxcr4b immunohisto-labeling and quantification of Barhl1a:GFP cells under the supervision of S.A. and L.P. All authors revised and approved the final version of the manuscript.

## Competing Interests

The authors declare no competing interests.

## Description of Additional Supplementary Files

### Supplementary Figure Legends

**Supplementary Figure 1.** Co-expression analysis of *lmx1b1* (red) and *barhl1a* (green) by double fluorescent *in situ* hybridization on 20 hours post-fertilization (hpf) embryos. Co-expression in the posterior tuberculum appears as yellow color (right panel). Scale bar: 20 *µ*m.

### Supplementary Movie Legends

**Supplementary Movie 1. Barhl1a:GFP expression in the retina recapitulates the wave of RGC genesis.** Time-lapse confocal imaging through the retina of a *Tg(barhl1a:GFP)* embryo from about 28 hpf. All time points are z-projections of confocal stacks imaged in lateral view. The retina is facing the eye, anterior is on the left, dorsal is at the top.

**Supplementary Movie 2. Barhl1a:GFP and Atoh7:gap43-RFP label retinal ganglion cells and their retinotectal projections.** 3D rendering and turn-around of the brain of a 35 hpf *Tg(barhl1a:GFP;atoh7:gap43-RFP)* embryo. The arrow points at the Barhl1a:GFP tectal projections.

**Supplementary Movie 3. Barhl1a expression in the diencephalon at 22 hpf.** 3D rendering and turn-around of the brain of a 22 hpf *Tg(barhl1a:GFP)* embryo

**Supplementary Movie 4. Expression domains of Barhl1a:GFP in the brain of a 35 hpf embryo.** Animation through the focus planes of a z-series stack of an *Tg(barhl1a:GFP;atoh7:gap43-RFP)* embryo in frontal view. Dt, dorsal thalamus; oc, optic chiasm; POC, post-optic commissure; vt, ventral thalamus; ot, optic tectum; pt, posterior tuberculum; hy, hypothalamus; pit, pituitary.

**Supplementary Movie 5. Barhl1a:GFP neurons extend commissural fibers in the ventral diencephalon.** Time-lapse confocal imaging through the brain of a *Tg(Barhl1a:GFP)* embryo from around 28 hpf. All time points are z-projections of confocal stacks imaged in frontal view.

**Supplementary Movie 6. Barhl1a:GFP expression highlights the formation of three commissures in the brain.** Time-lapse confocal imaging through the brain of a *Tg(barhl1a:GFP)* embryo from around 28 hpf in lateral view. Anterior is on the left, dorsal is up. All time points are z-projections of confocal stacks imaged in lateral view (retina facing the eye, anterior is on the left, dorsal is at the top). The eye is visible only in the first time-point. POC, post-optic commissure; OC, Optic chiasm; PC, posterior commissure.

**Supplementary Movie 7. Mosaic expression of Barhl1a:gal4/UAS:RFP cells showing their fiber extension along the post-optic commissure.** Frontal animation through the focus planes of a z-series stack of an *Tg*(*UAS:RFP;cry:GFP*) embryo injected at 1-cell stage with the *barhl1.2:gal4-vp16* BAC construct at around 28 hours post-fertilization. Red arrows show single RFP cells projecting their fibers ventrally along the optic tract in the basal diencephalon and single cells extending their axons along the PC in the presumptive ventral midline.

